# Exposing hidden alternative backbone conformations in X-ray crystallography using qFit

**DOI:** 10.1101/018879

**Authors:** Daniel A. Keedy, James S. Fraser, Henry van den Bedem

**Affiliations:** Department of Bioengineering and Therapeutic Sciences, University of California, San Francisco, San Francisco, California, United States of America; Division of Biosciences, SLAC National Accelerator Laboratory, Stanford University, California, United States of America

## Abstract

Proteins must move between different conformations of their native ensemble to perform their functions. Crystal structures obtained from high-resolution X-ray diffraction data reflect this heterogeneity as a spatial and temporal conformational average. Although movement between natively populated alternative conformations can be critical for characterizing molecular mechanisms, it is challenging to identify these conformations within electron density maps. Alternative side chain conformations are generally well separated into distinct rotameric conformations, but alternative backbone conformations can overlap at several atomic positions. Our model building program qFit uses mixed integer quadratic programming (MIQP) to evaluate an extremely large number of combinations of sidechain conformers and backbone fragments to locally explain the electron density. Here, we describe two major modeling enhancements to qFit: peptide flips and alternative glycine conformations. We find that peptide flips fall into four stereotypical clusters and are enriched in glycine residues at the *n*+1 position. The potential for insights uncovered by new peptide flips and glycine conformations is exemplified by HIV protease, where different inhibitors are associated with peptide flips in the “flap” regions adjacent to the inhibitor binding site. Our results paint a picture of peptide flips as conformational switches, often enabled by glycine flexibility, that result in dramatic local rearrangements. Our results furthermore demonstrate the power of large-scale computational analysis to provide new insights into conformational heterogeneity. Overall, improved modeling of backbone heterogeneity with high-resolution X-ray data will connect dynamics to the structure-function relationship and help drive new design strategies for inhibitors of biomedically important systems.

**Author Summary:** Describing the multiple conformations of proteins is important for understanding the relationship between molecular flexibility and function. However, most methods for interpreting data from X-ray crystallography focus on building a single structure of the protein, which limits the potential for biological insights. Here we introduce an improved algorithm for using crystallographic data to model these multiple conformations that addresses two previously overlooked types of protein backbone flexibility: peptide flips and glycine movements. The method successfully models known examples of these types of multiple conformations, and also identifies new cases that were previously unrecognized but are well supported by the experimental data. For example, we discover glycine-driven peptide flips in the inhibitor-gating “flaps” of the drug target HIV protease that were not modeled in the original structures. Automatically modeling “hidden” multiple conformations of proteins using our algorithm may help drive biomedically relevant insights in structural biology pertaining to, e.g., drug discovery for HIV-1 protease and other therapeutic targets.

## Introduction

Even well-folded globular proteins exhibit significant flexibility in their native state [1]. However, despite advances in nuclear magnetic resonance dynamics experiments and computational simulations, accurately characterizing the nature and extent of biomolecular flexibility remains a formidable challenge [2]. While traditionally X-ray crystallography is associated with characterizing the ground state of a biomolecule, the ensemble nature of diffraction experiments means that precise details of alternative conformations can be accessed when the electron density maps are of sufficient quality and resolution [3]. These maps represent spatiotemporal averaged electron density from conformational heterogeneity across the millions of unit cells within a crystal [4, 5].

Computational methods have made strides toward uncovering and modeling conformational heterogeneity in protein structures from crystallographic data [3]. However, there is currently no automated approach to recognize the features of extensive backbone flexibility in electron density maps, model the constituent alternative conformations, and validate that the incorporation of heterogeneity improves the model. B-factors theoretically model harmonic displacements from the mean position of each atom, but in practice are often convolved with occupancies of discrete alternative positions when multiple backbone conformations partially overlap [5]. Statistical analyses of electron density using Ringer has revealed evidence for a surprising number of “hidden” alternative conformations in electron density maps [6, 7]. The phenix.ensemble_refinement method [8] uses electron density to bias molecular dynamics simulations, then assembles snapshots from this trajectory into a multi-copy ensemble model. However, energy barriers of the simulation may prevent sampling of well separated backbone conformations. Accurately modeling protein conformational heterogeneity, in particular when the mainchain adopts distinct conformations for one or a number of contiguous residues, remains a difficult task. The spatial overlap of electron density of multiple conformations and the relatively similar profiles of branching mainchain and sidechains blur structural features that can guide the human eye to reduce the large number of possible interpretations [9].

We have previously developed qFit [10], a method for automatically disentangling and modeling alternative conformations and their associated occupancies, which are represented by the variable *q* (for “oc*<u>cu</u>*pancy”) in standard structure factor equations. The qFit algorithm examines a vast number of alternative interpretations of the electron density map simultaneously. To propitiously explore a high-dimensional search space, conformational sampling is guided by the anisotropy of electron density at the Cβ atom position, the nexus of backbone and sidechain in polypeptides [11]. For each slightly shifted Cβ atom position, qFit samples sidechain conformations with a rotamer library [12] and uses inverse kinematics to maintain backbone closure [9]. Finally, it selects a set of one to four conformations for each residue that, collectively, optimally explain the local electron density in real space.

However, the anisotropy of the Cβ atom limits the exploration radius of qFit to model backbone conformational heterogeneity. While protein backbone motions are often associated with large-amplitude conformational flexibility of surface loop regions, subtle motions can have important ripple effects in closely packed areas via sidechain-backbone coupling. For example, fast (ps-ns) backbone NH and sidechain methyl order parameters from spin relaxation experiments are highly correlated with each other in flexible regions [13], suggesting that mainchain and sidechain motions collectively sample conformational substates. For example, a backbone backrub motion [14] repositions the Cα-Cβ bond vector in a plane perpendicular to the chain direction, enabling the sidechain to access alternative, often sparsely populated rotamers that otherwise would be energetically unfavorable. We previously linked coupled transitions between alternative sidechain conformations, like “falling dominos”, to enzymatic turnover and allostery [15, 16].

Additionally, qFit cannot model discrete conformational substates such as peptide flips, which are >90° rotations of a peptide group while minimally perturbing the flanking residues. Some structure validation methods highlight incorrect peptide orientations [17] and even automate subsequent model rebuilding [18]. However, rebuilding fits a correct, unique conformation rather than multiple well-populated alternative peptide conformations. Peptide flips can have important functional roles in proteins. For example, flavodoxin undergoes peptide rotations between functional states as part of the catalytic cycle [19], and peptide flips that convert β-sheet to α-sheet have been linked to amyloid formation [20]. Furthermore, high-resolution crystal structures have shown that alternative conformations related by a peptide flip may be populated in the same crystal [14].

Modeling alternative conformations of glycine residues, which lack a Cβ atom, is also a current limitation of qFit. The lack of a Cβ atom allows glycine residues to access otherwise forbidden regions of conformational space [11] and thereby fill special structural roles such as capping helix C-termini [21]. In addition, the flexibility of glycines may contribute directly to function at flexible inter-domain linkers or conformationally dynamic enzyme active sites [22]. Automatically modeling such cases as alternative conformations with qFit paves the way toward understanding their contributions to protein function. Increasingly, new experiments are being proposed which, combined with computational analysis, can extract the spatiotemporal ensemble from electron density maps [15, 23, 24].

Adding the capability to model peptide flips and alternative conformations for glycines will increase our power to uncover conformational heterogeneity. While the number of sampled conformations for glycines is modest owing to a missing side-chain, including peptide flips for all amino acids adds significant computational complexity to the qFit algorithm. A powerful quadratic programming algorithm lies at the core of qFit and is necessary to determine non-zero occupancies for up to four conformations from among hundreds or even thousands of candidate conformations for each residue. Even for modest sample sizes, around 500, the number of combinations of candidate conformations is enormous, exceeding 10^9^. As more backbone motion is incorporated into qFit, the computational complexity increases, demanding a parallelized approach to refinement on a residue by residue basis. Although this moves rebuilding away from a single node towards a larger compute cluster, the combination of data-driven sampling and selection has enabled qFit to automatically build multiconformer models that have illuminated intramolecular networks of coupled conformational substates [16] and the effects of cryocooling crystals [25, 26]. Similar hybrid approaches using robotics sampling and selection based on experimental NMR data are also being extended to nucleotide systems such as the excited state of HIV-1 TAR RNA [27].

Here we introduce qFit 2.0, an updated version of the qFit algorithm with new capabilities for modeling near-native backbone conformational heterogeneity in crystal structures. We first describe the quadratic programming procedure that allows selection of a small set of conformations per residue that collectively account for the local electron density, and discuss its extension to fitting backbone atoms in addition to sidechain atoms. We then describe new conformational sampling features of qFit 2.0, in particular glycine shifts and peptide flips. Finally, we validate the updated algorithm with both synthetic and experimental X-ray data. qFit 2.0 is freely available by webserver and source code is available for download at https://simtk.org/home/qfit.

## Results

### Improved backbone sampling and selection in qFit

To automatically identify alternative backbone conformations, including peptide flips, we augmented the sample-and-select protocol in qFit (see Figure 1 and Methods). Previously, conformations were sampled based on anisotropy of the Cβ atom and were selected based on the fit between observed and calculated electron density for the sidechain (Cβ atom and beyond) only. Alternative conformations for mainchain atoms were ultimately included in the multiconformer model only because they accommodated the best sidechain fits. In qFit 2.0, we now select conformations based on the fit between observed and calculated electron density for the sidechain atoms and also the backbone O atom. The O atom is an excellent yardstick for identifying backbone conformational heterogeneity for two reasons. First, it is furthest from the Cα-Cα axis so its density profile is somewhat isolated and is displaced most by rotations around that axis [14]. Second, it has more electrons than other backbone heavy atoms, so is most evident in electron density maps. This change allows us to select peptide flips outside of α-helices and β-sheets, where flips are prevented by steric and hydrogen-bonding constraints, then directly select flipped conformations. This procedure is effective because the large movement of the backbone O during a peptide flip leaves a major signature in the electron density.

**Figure 1:**
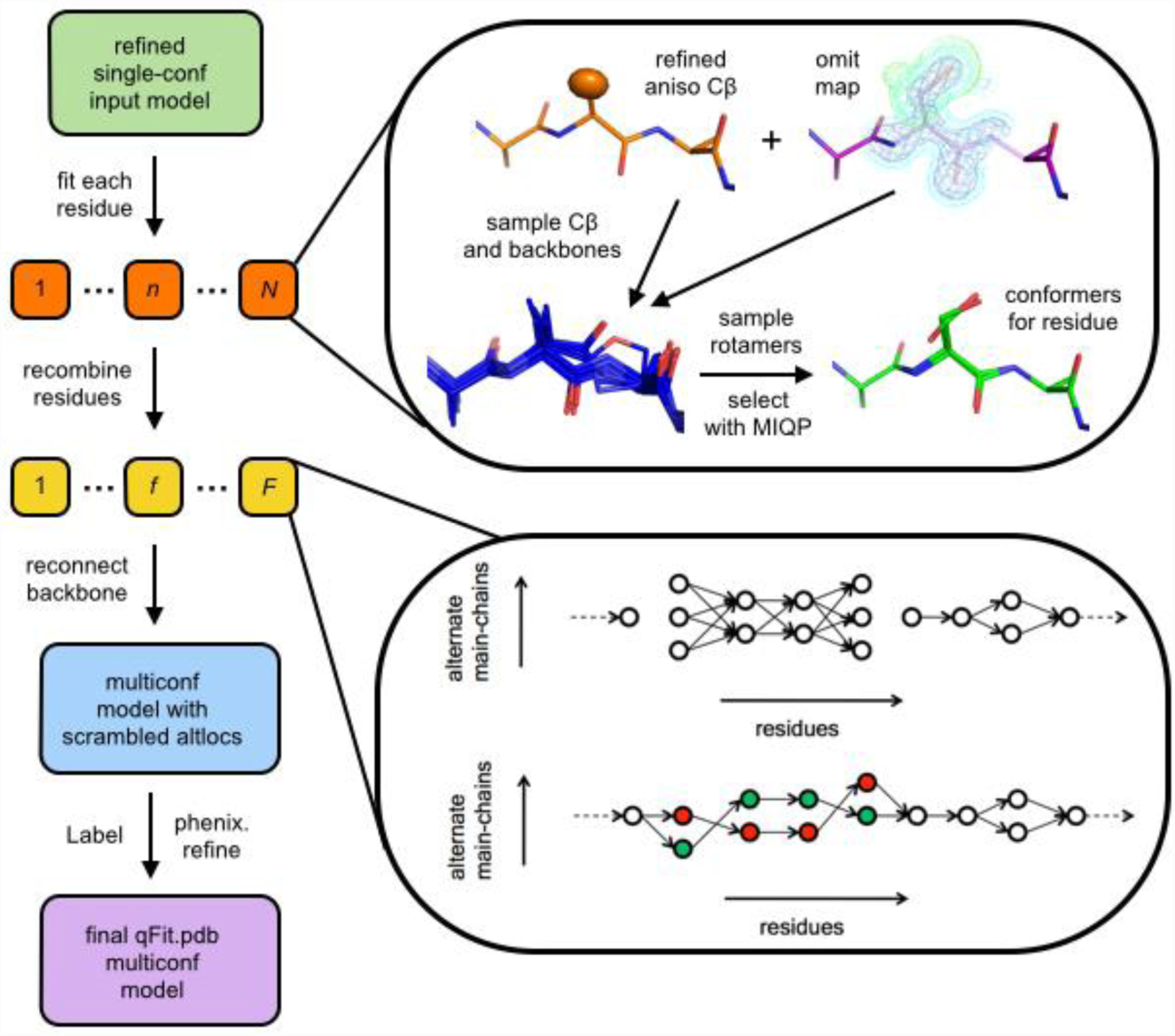
Flowchart of the qFit 2.0 algorithm. qFit can operate on each residue in the protein (orange boxes) in parallel (1 ≤ *n* ≤ *N* indices are for residues in the protein). Anisotropic refinement gives a thermal ellipsoid for the Cβ (orange model), and refinement with occupancies set to 0 gives an omit map (purple model). These inputs are combined, backbone translations and peptide flips are sampled (blue models), each backbone is decorated with sidechain rotamers, and an MIQP is used to select 1-4 conformations for the residue. Residues with consecutive multiple backbone conformations, called fragments (yellow boxes), are then subjected to a second MIQP to trace compatible alternative backbone conformations across residues. Residues and fragments are combined into an intermediate model. Finally, a Monte Carlo procedure is used to adjust alternative conformation labels (“altloc” identifiers) to minimize steric overlaps, and the final model is refined.

### Glycine modeling

Incorporating the backbone O atom also enhances the detection of less discrete backbone conformational changes. In particular, we now sample alternative glycine conformations based on anisotropy of the electron density for the O atom, by analogy to the Cβ-driven sampling for all other amino acids. This results in alternative glycine conformations that are dictated by their own local electron density. After sampling, we select combinations of conformers from a pool of candidates based on both sidechain and backbone O atoms for all amino acids, including glycines. This addition results in greater potential to discover alternative conformations throughout the protein and include additional conformational heterogeneity in the final multiconformer model.

### Characterizing peptide flip geometry

The nullspace inverse kinematics procedure of qFit [9] naturally encodes backrub [14], crankshaft [28], and shear [29, 30] motions (**Figure S1**) where they are dictated by the anisotropy of the electron density for the Cβ atom. However, this anisotropy cannot identify more discrete substates of the backbone, such as peptide flips. Peptide flips are large, ∼180° rotations of a peptide plane in protein backbone with minimal disturbance of adjacent peptide conformations. Enumerating many peptide flip candidate conformations with the nullspace inverse kinematics procedure would quickly lead to prohibitively large sample sizes. We therefore examined common geometries of discrete peptide flips to expedite sampling of discrete backbone substates in qFit 2.0.

Steric interactions prevent arbitrary rotations of the peptide plane, much like sidechains adopt preferred rotamer conformations. To identify plausible geometries for peptides relative to a single input peptide, we examined cases where the peptide rotates by 90-180° around the Cα-Cα axis. We identified 147 peptide flips modeled as alternative conformations in high-quality structures. After filtering this set of peptide flips with structure validation criteria and reserving some examples for a test set, we retained 79 examples that clustered around four geometries (**Table S1**). We observed that peptide flips often included rotation and translation within the peptide plane such that the first Cα moves “below” the Cα-Cα axis and the second Cα moves “above” it (from the view in Figure 2A,C). These in-plane movements justify sampling geometries found in natural peptide flips in qFit 2.0 rather than, e.g., simply rotating the peptide 180° around the Cα-Cα axis. The first two clusters, “simple down” (Figure 2A,C, blue) and “tweaked down” (Figure 2A,C, red), feature a very nearly 180° rotation around the Cα-Cα axis, but with different in-plane adjustments. By contrast, the second two clusters, “left” (Figure 2B,D, green) and “right” (Figure 2B,D, brown), feature rotations closer to 120°, but in opposite directions. Our dataset here is sufficient to propose plausible, well-validated peptide flip geometries for sampling in qFit 2.0, and suggests that the four clusters could also be used to inspire moves in protein design.

**Figure 2:**
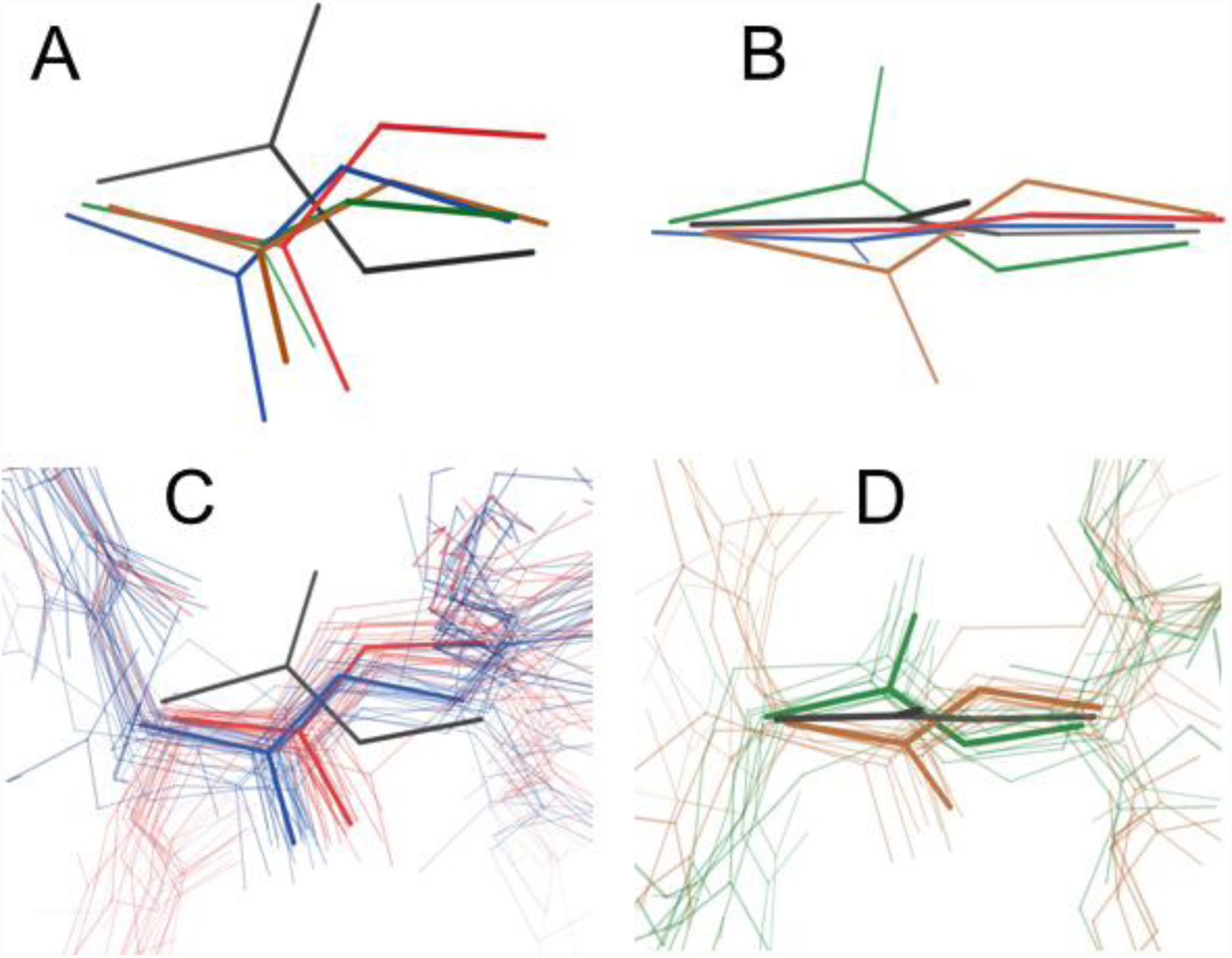
Geometry and distribution of peptide flips in training set. **(A,B)** Reference primary conformation peptide (black) and four cluster centroids for secondary peptide conformations (colors), from the side **(A)** or “top-down” **(B)**. **(C)** Members from the training set segregate into two ∼180° rotated clusters with different translations in the peptide plan (blue vs. red). View from roughly the same angle as **(A)**. **(D)** Other members from the training set segregate into +120° and -120° rotated clusters (green vs. brown). View from roughly the same angle as **(B)**.

### Structural context of flips

We found that the two “down” clusters were more common in tight turns between β-strands: 41-50% of flips in these clusters were found in turns, as compared to 0-14% for the other two flip clusters (with a conservative definition of a turn; see Methods) (**Table 1**). The flip is nearly always associated with a transition between Type I/I’ and II/II’ turns. The “left”/”right” clusters were dispersed among many irregular structural contexts, but not α-helices or β-sheets. Across the four clusters, the first residue of the peptide was a glycine 7.5% of the time, in line with the general abundance of glycines in proteins (7-8%). However, the second residue of the peptide was a glycine significantly more frequently (50%, p < 10^-22^). This was true for the “left”/”right” clusters (21%, p < 0.05) and especially the two “down” clusters (Figure 2C) (64%, p < 10^-24^). This may be in part because a glycine as the second residue of a peptide can lower the flip transition energy [31]. These results generally agree with reports of flip-like conformational differences between the same tight turn in separate homologous structures [32].

**Table 1:** Peptide flip geometries aggregate into distinct clusters. Colors refer to Figure 2.

| Cluster | # of examples | # (%) in tight turn | # (%) with Gly as first residue | # (%) with Gly as second residue |
| --- | --- | --- | --- | --- |
| “tweaked down” (red) | 26 | 13 (50%) | 0 (0%) | 16 (62%) |
| “simple down” (blue) | 29 | 12 (41%) | 4 (14%) | 19 (66%) |
| “left” (green) | 10 | 0 (0%) | 1 (10%) | 2 (20%) |
| “right” (brown) | 14 | 2 (14%) | 1 (7%) | 3 (21%) |

### Tests with synthetic datasets

To test these advances, we first explored synthetic datasets spanning resolutions from 0.9 to 2.0 Å with increasing B-factors as a function of resolution and Gaussian noise added to structure factors (see Methods). We used the Top8000 peptide flip geometry cluster centroids, with the alternative conformations at 70/30 occupancies for the “tweaked down” cluster and 50/50 occupancies for the other three clusters. Because qFit uses these geometries to sample peptide flips, we expected it would be able to successfully identify each flipped alternative conformation starting from the primary (labeled “A”) conformation at high-to-medium simulated resolution, but less well at lower simulated resolution. Indeed, qFit 2.0 successfully finds the flipped conformations for most peptide flip geometry clusters across resolutions with a 92% success rate overall; this rate drops only slightly with resolution from 0.9 to 2.0 Å (Figure 3). Since we rebuilt the entire protein chain, we also assessed the performance on other residues. By contrast to the true positive peptide flip results, the peptide flip and rotamer false positive rates remain quite low across clusters and resolutions (Figure 3). These results indicate that qFit 2.0 is effective at identifying peptide flip alternative conformations across a wide range of crystallographic resolutions without introducing spurious conformations.

**Figure 3:**
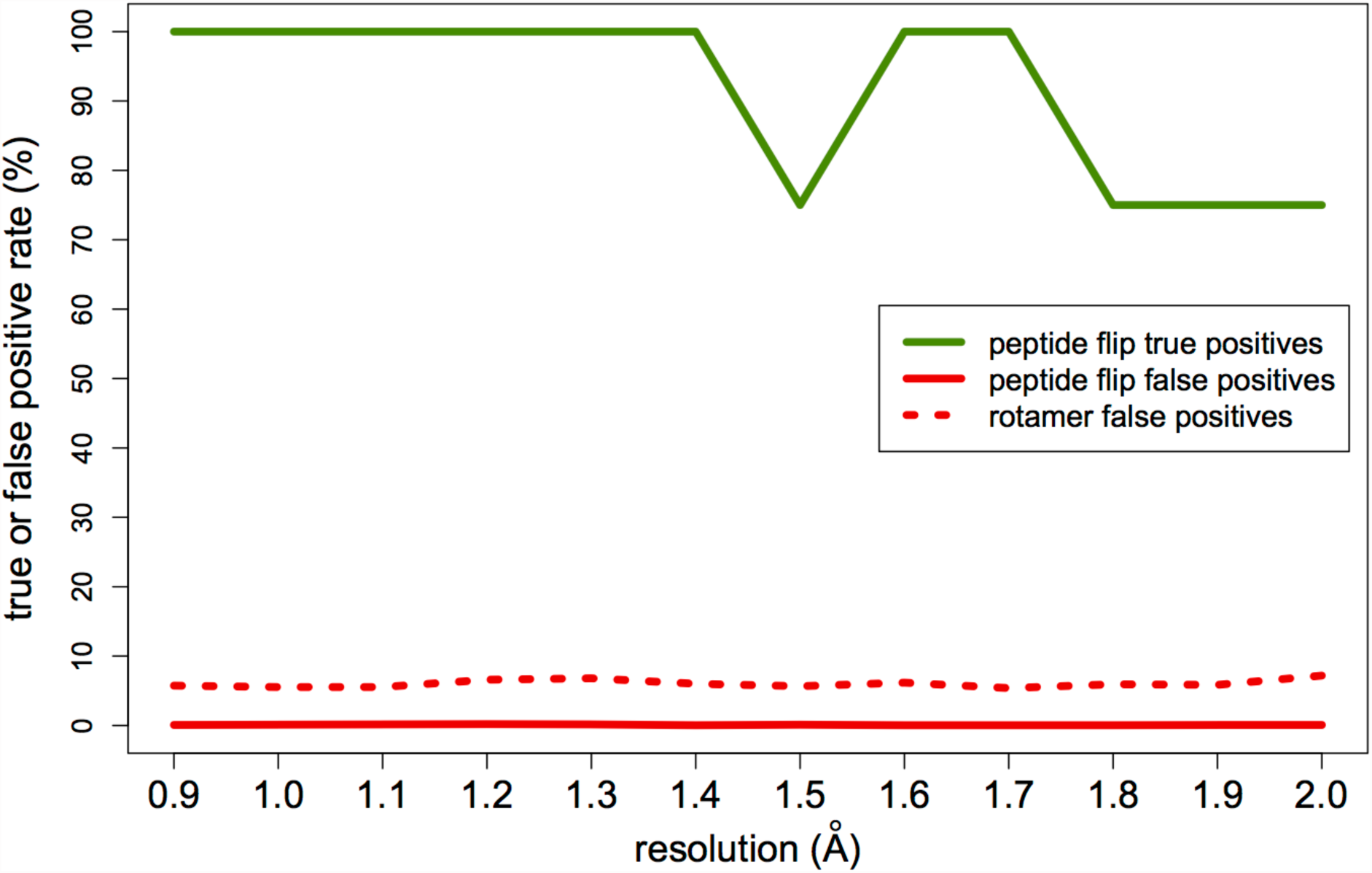
True vs. false positives with synthetic data. Peptide flip true positives = percent of peptide flips in the actual synthetic model that are present in the qFit 2.0 model. Peptide flip false positives = percent of residues with a peptide flip in the qFit 2.0 model that are not in the actual synthetic model. Rotamer false positives = percent of sidechain rotamers (as defined by MolProbity [12, 33]) in the qFit 2.0 model that are not in the actual synthetic model. True positives in green; false positives in red. Peptide flips in solid lines; rotamers in dotted line. Data is averaged over all four synthetic datasets (corresponding to the four peptide flip geometry clusters in Figure 2) and all three mainchain amplitudes are considered; see Methods.

### Tests with experimental datasets

Although tests with synthetic datasets offer insight into resolution dependence, a more direct test of the usefulness of qFit 2.0 involves crystal structures with real data. We combined structures left out of the training set from the Top8000 peptide flip examples with a few more manually curated examples for a total of 15 test cases (Table 2). When comparing qFit 2.0 models to rerefined original structures, R__free__ is better for 7/15 cases and R_work_ is better for 8/15 cases (**Figure S2**). However, after rerefinement with automated removal and addition of water molecules to allow the ordered solvent to respond to the new protein alternative conformations modeled by qFit (see Methods), R_free_ is better for the qFit 2.0 model for 10/15 cases and R_work_ is better for 13/15 cases (Figure 4). The differences generally are small: the average ΔR_free_ is ∼0.1%. Overall, these results suggest that qFit 2.0 models explain experimental crystallographic data as well as or better than traditional refinement protocols at a global structural level.

**Figure 4:**
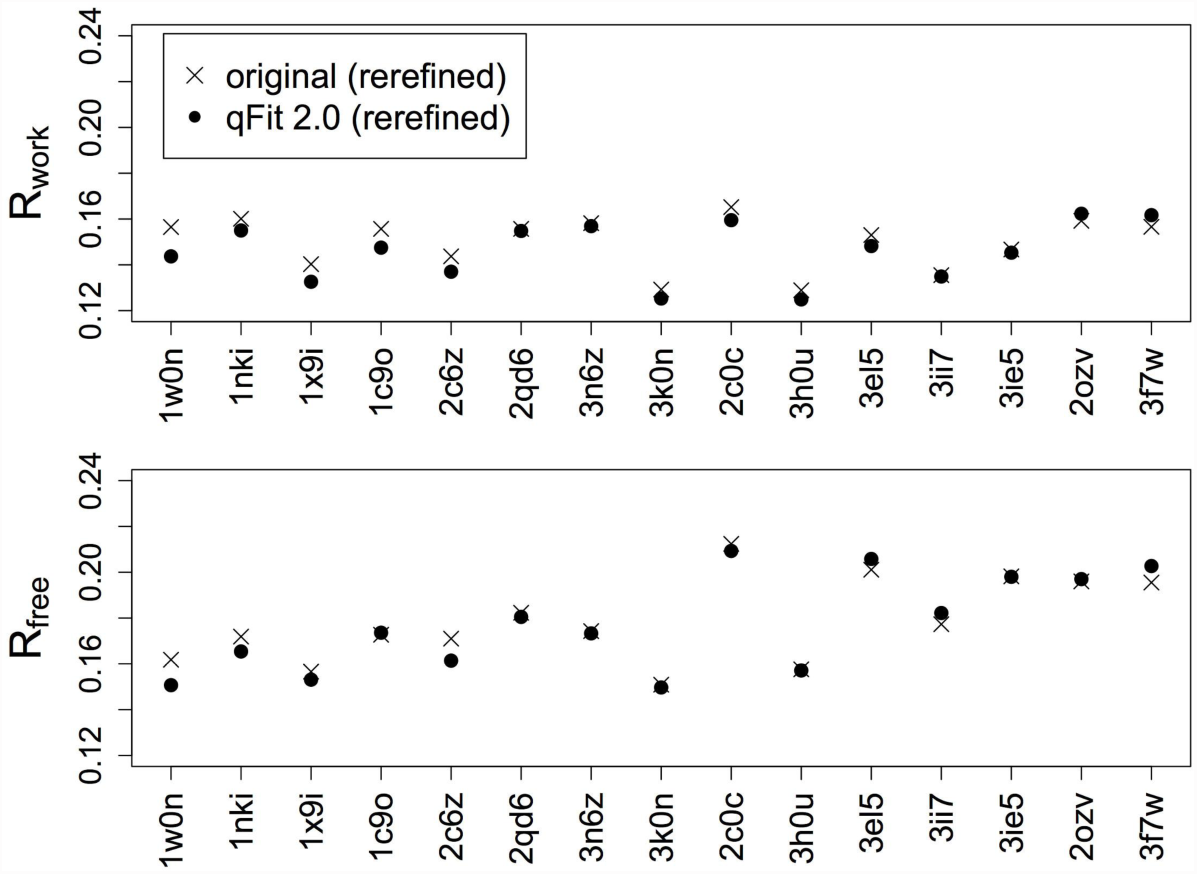
Multiconformer modeling with qFit results in similar or better crystallographic R-factors. R_work_ and R_free_ are plotted vs. PDB ID sorted from high to low resolution. X’s indicate rerefined original structures and filled circles indicate qFit 2.0 models; both are after refinement with water picking.

**Table 2:** List of positive-control peptide flip test cases. Last column indicates whether or not qFit 2.0 found the peptide flip alternative conformations for at least one of the three backbone amplitude parameters. Overall, 14/18 (78%) peptide flips were successfully identified.

| PDB ID | Resolution (Å) | Chain | Peptide | Found flip? |
| --- | --- | --- | --- | --- |
| 1w0n | 0.80 | A | 42-43 | Y |
| 1nki | 0.95 | A | 53-54 | Y |
| 1nki | 0.95 | B | 53-54 | Y |
| 1x9i | 1.16 | A | 65-66 | Y |
| 1c9o | 1.17 | A | 36-37 | Y |
| 1c9o | 1.17 | B | 36-37 | n |
| 2c6z | 1.20 | A | 227-228 | n |
| 2qd6 | 1.28 | A | 50-51 | n |
| 2qd6 | 1.28 | B | 150-151 | Y |
| 3n6z | 1.30 | A | 177-178 | Y |
| 2c0c | 1.45 | A | 44-45 | n |
| 3h0u | 1.50 | C | 115-116 | Y |
| 3el5 | 1.60 | A | 50-51 | Y |
| 3el5 | 1.60 | B | 50-51 | Y |
| 3ii7 | 1.63 | A | 539-540 | Y |
| 3ie5 | 1.69 | A | 61-62 | Y |
| 2ozv | 1.70 | A | 54-55 | Y |
| 3f7w | 1.85 | A | 48-49 | Y |

While global metrics are important, a major focus of the current work is correctly identifying local alternative backbone conformations. To explore this aspect, we compared results from qFit 2.0 to those from qFit 1.0 and original deposited structures for our test set (Table 2). qFit 2.0 successfully models both flipped conformations in 14/18 (78%) cases. For example, Val539-Gly540 in the Kelch domain of human KLHL7 is modeled with two alternative conformations related by a peptide flip (1.63 Å, PDB ID 3ii7) (Figure 5A). qFit 1.0 fails to discover the flip, resulting in significant difference electron density peaks (Figure 5B). By contrast, qFit 2.0 beautifully recovers both alternative conformations (Figure 5C). In another example, Asn42-Gly43 in carbohydrate binding domain 36 at high resolution (0.8 Å, PDB 1w0n) adopts flipped peptide conformations - yet MolProbity flags geometry errors in the deposited structure that indicate it re-converges too quickly, with alternative conformations for only the Asn42 and not also Gly43 (Figure 5D). qFit 1.0 fails to capture the flip (Figure 5E). However, qFit 2.0 not only identifies both peptide flip conformations for Asn42, but also includes split conformations for Gly43, thereby repairing the covalent backbone geometry (Figure 5F). In both cases, the peptide flip and glycine sampling enhancements in qFit 2.0 combine to model discrete backbone heterogeneity as accurately as or even better than the original structure.

**Figure 5:**
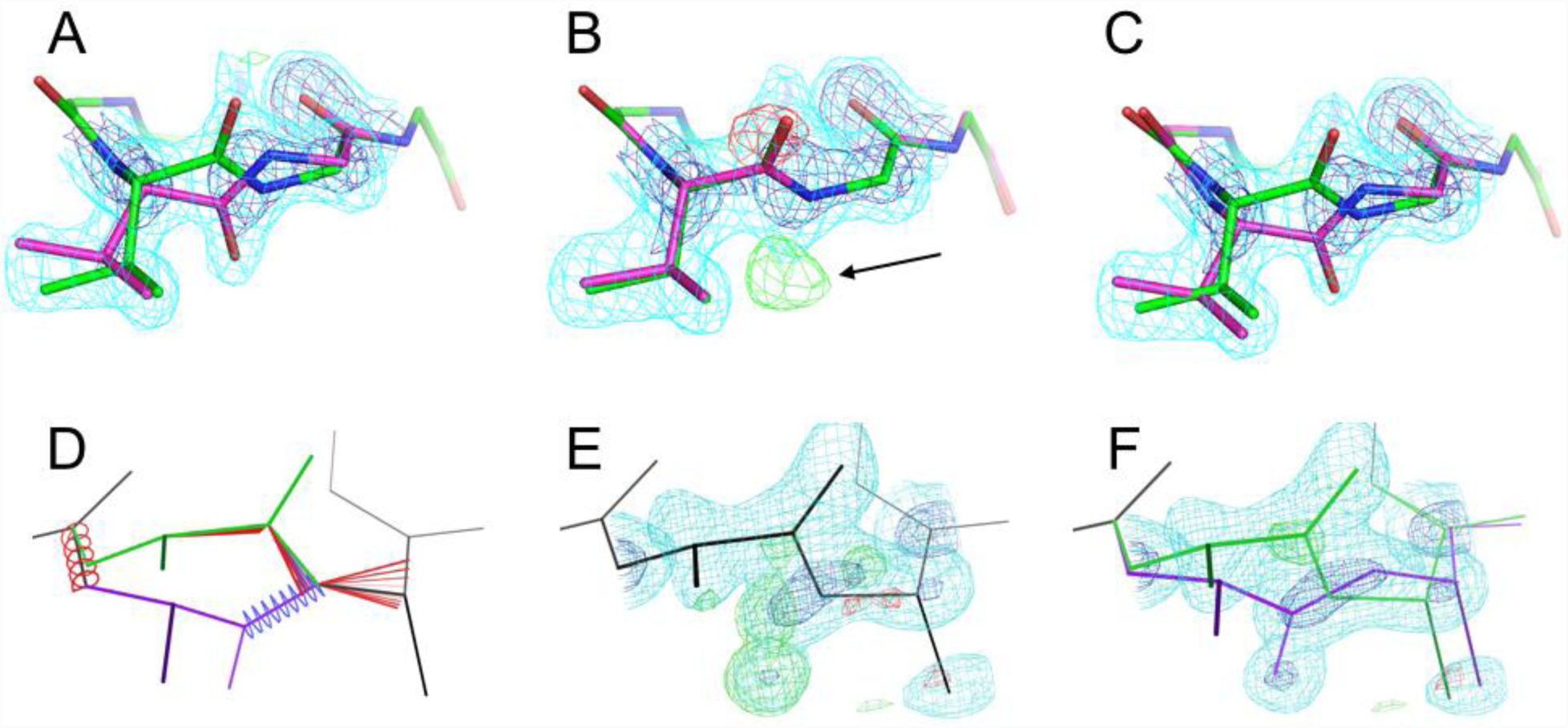
qFit 2.0 successfully identifies known peptide flips. **(A-C)** Val539-Gly540 in the Kelch domain of human KLHL7 at 1.63 Å (PDB ID 3ii7). 2mFo-DFc electron density is contoured at 1.2 σ (cyan) and 2.5 σ (blue); mFo-DFc electron density is contoured at +3.0 σ (green) and -3.0 σ (red). **(A)** The deposited model includes alternative conformations for this peptide, which are well justified by the electron density. **(B)** qFit 1.0 starting from single-conformer input fails to find the second conformation, resulting in peaks in the difference density map (arrow). **(C)** qFit 2.0 finds both conformations, resulting in the disappearance of the difference peaks. **(D-E)** Asn42-Gly43 in carbohydrate binding domain 36 at 0.8 Å resolution (PDB ID 1w0n). The Asn42 sidechain (left, darker green/purple) points up out of the image so is visually truncated. In **(E-F)**, 2mFo-DFc electron density is contoured at 1.5 σ (cyan) and 2.5 σ (blue); mFo-DFc electron density is contoured at +3.0 σ (green) and -3.0 σ (red). **(D)** The deposited structure includes alternative conformations (green and purple) related by a peptide flip, but re-converges too early at the Gly43 backbone N atom, resulting in >4 σ bond length (red and blue fans) and bond angle (red and blue springs) outliers [33]. **(E)** qFit 1.0 fails to identify the flip, leaving significant difference density map features. **(F)** qFit 2.0 identifies the flip at Asn43 and also correctly splits Gly43 into separate conformations, thereby flattening the difference map relative to qFit 1.0 and eliminating the covalent geometry errors in the original structure.

### Discovering new conformational heterogeneity from experimental datasets

In addition to retrospective positive-control tests, we also looked prospectively for “hidden” peptide flip alternative conformations that are unmodeled in existing structures. One such example is Met519-Thr520 in RNA binding protein 39. In chain A of the room-temperature structure (PDB ID 4j5o), the mFo-DFc difference electron density map around this peptide has significant positive and negative peaks, indicating it is mismodeled as a single conformation (Figure 6A). Other instances of this peptide – including in chain B of the room-temperature structure and both chains of the cryogenic structure – feature conformational diversity, much of which may be related to crystal contacts; however, these conformations fail to account for the room-temperature chain A mFo-DFc peaks (Figure 6B). However, using the room-temperature data, qFit 2.0 identifies a peptide flip in this region, which repositions Met519 and flattens the local difference density (Figure 6C). By contrast, it does not identify a peptide flip for this region in either chain using the cryogenic data, which is in accord with previous reports that cryocooling crystals can conceal or otherwise perturb conformational heterogeneity that is present at room temperature [25, 26].

**Figure 6:**
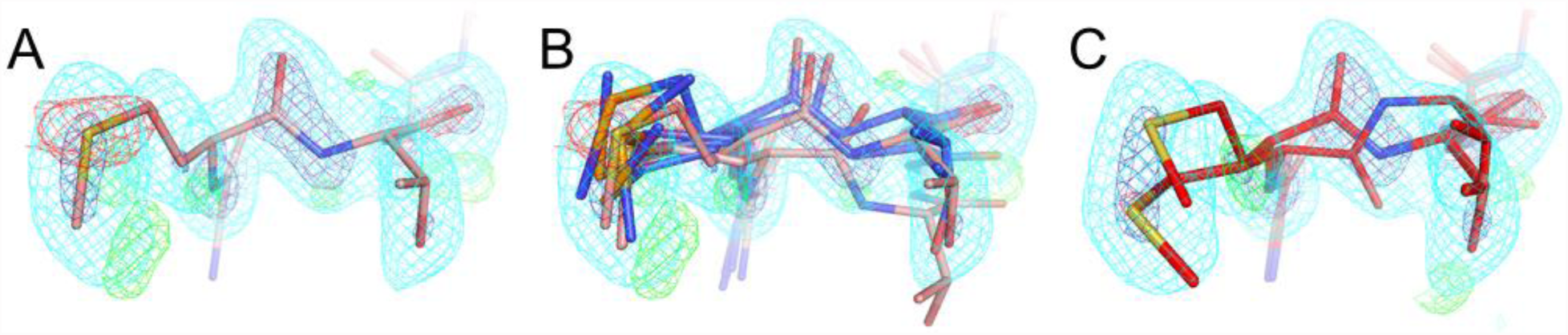
qFit 2.0 finds a hidden peptide flip at room temperature. **(A)** Met519-Thr520 in mouse RNA binding protein 39 (RBM39) is modeled with just a single conformation in chain A of the 1.11 Å room-temperature structure (PDB ID 4j5o, pink). There appear to missing unmodeled conformations based on mFo-DFc difference electron density contoured at +3.0 σ (green) and -3.0 σ (red). 2mFo-DFc electron density is shown contoured at 0.9 σ (cyan) and 2.5 σ (dark blue). **(B)** Although there is diversity for this region in chain B of the asymmetric unit from this structure and in chains A and B from the 0.95 Å cryogenic structure (PDB ID 3s6e, blue), none of the other instances explain the electron density at RT in chain A. There is also no clear evidence for missing alternative conformations in these other instances (not shown). **(C)** The RT qFit model (magenta) adds a flipped peptide as an alternative conformation, which positions the Met519 sidechain differently. Collectively, these changes better explain the local electron density.

In addition to selection of conformers based on fit to density for the backbone O atom for all amino acids, qFit 2.0 also adds sampling based on this atom for glycine, enabling density-driven backbone sampling for the most flexible amino acid. This facilitates modeling peptide flips in which one of the constituent residues is a glycine, as seen in the examples above (Figure 5) -- but also opens the door to modeling less discrete glycine flexibility. For the 489 glycines across the 15 datasets in the test set (Table 2), qFit 1.0 cannot model more than a single conformation, but qFit 2.0 models alternative conformations for 365/489 (75%) of glycines. The Cα displacements average 0.28 Å and range from <0.01 Å up to 1.70 Å. Only 4 (4%) of these glycines were modeled with alternative conformations in the original PDB structures. These results show that the direct sampling and selection based on electron density for glycine backbone atoms in qFit 2.0 successfully identify conformational heterogeneity that was formerly unrecognized. For example, a small, glycine-rich loop in PDB ID 3ie5 is modeled with a single conformation in the deposited structure and qFit 1.0 model (Figure 7A). By contrast, qFit 2.0 recognizes the anisotropy of the electron density for each of the three glycine O atoms in the loop, so models them with alternative conformations that collectively shift the entire mini-loop region (Figure 7B).

**Figure 7:**
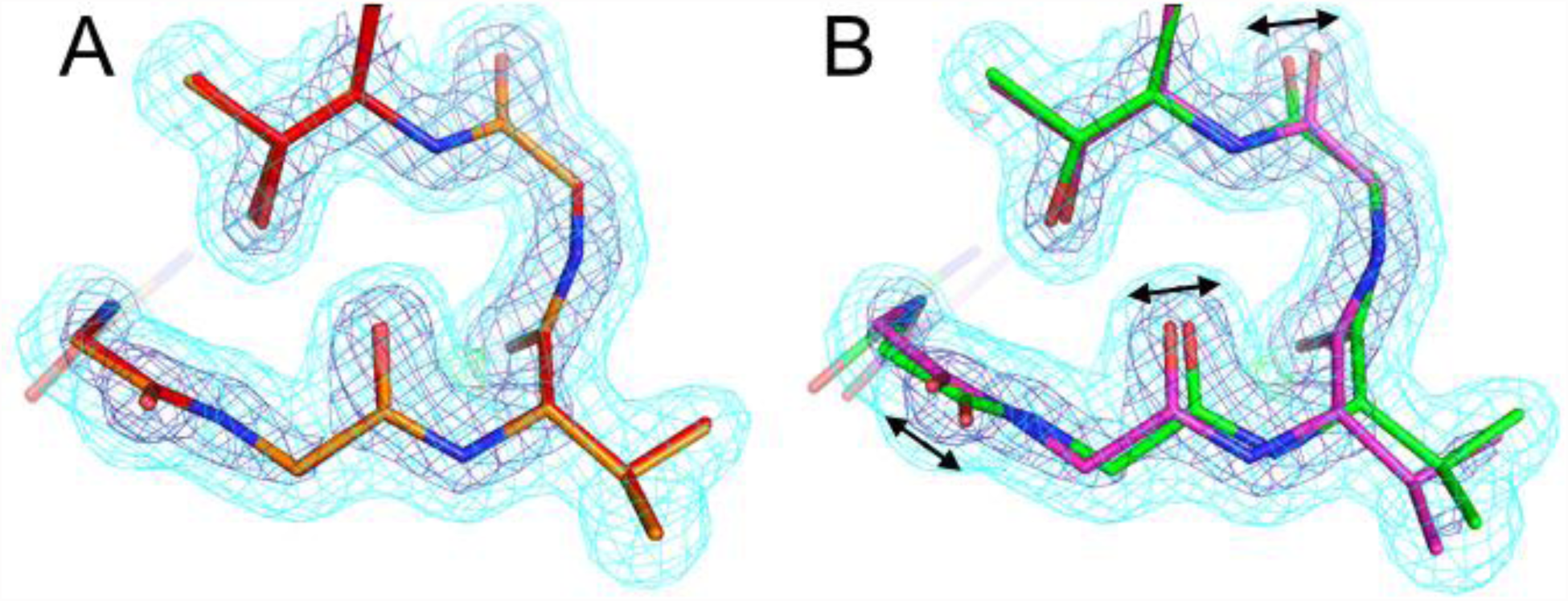
qFit 2.0 identifies alternative glycine conformations. This small loop in the 1.69 Å structure of Hyp-1 protein from St. John’s wort (PDB ID 3ie5) includes several glycines: 49, 50, and 52. **(A)** The deposited structure (orange) depicts these glycines with single conformations. The qFit 1.0 model (red) does the same, because it cannot sample alternative glycine conformations. **(B)** The qFit 2.0 model identifies alternative conformations (green/purple) for the entire loop, including all three glycines, based on subtly anisotropic backbone O atoms (arrows). 2mFo-DFc electron density contoured at 1.0 σ (cyan) and 3.0 σ (blue); mFo-DFc electron density contoured at +3.0 σ (green) and -3.0 σ (red).

Selecting conformers based on fit to density for the backbone O atom helps find alternative conformations not only for glycines, but also more generally for other amino acids. In many cases, this additional data-driven aspect to conformer selection drives the identification of subtle, non-discrete backbone motions that are coupled to larger, discrete sidechain changes. Indeed, for the 15 proteins in Table 2, qFit 2.0 shifts the Cα more than does qFit 1.0 for 52% of residues, but the reverse is true for only 20% of residues (the remaining residues are not moved by either version) (Figure 8A). Furthermore, for 63% of the residues for which qFit 2.0 finds a new sidechain rotamer that qFit 1.0 does not, qFit 2.0 also moves the Cα more (Figure 8B). These results imply that the backbone sampling by qFit 2.0 not only increases backbone heterogeneity in and of itself, but also drives discovery of sidechain conformational heterogeneity. As one specific example, Thr157 in cyclophilin A is modeled with alternative backbone and rotamer conformations in the deposited structure (Figure 8A). qFit 1.0 fails to find the alternative rotamer because it maintains a single backbone conformation (Figure 8B), but, driven by carbonyl O anisotropy, qFit 2.0 identifies the alternative backbone conformations, allowing it to discover the second rotamer (Figure 8C).

**Figure 8:**
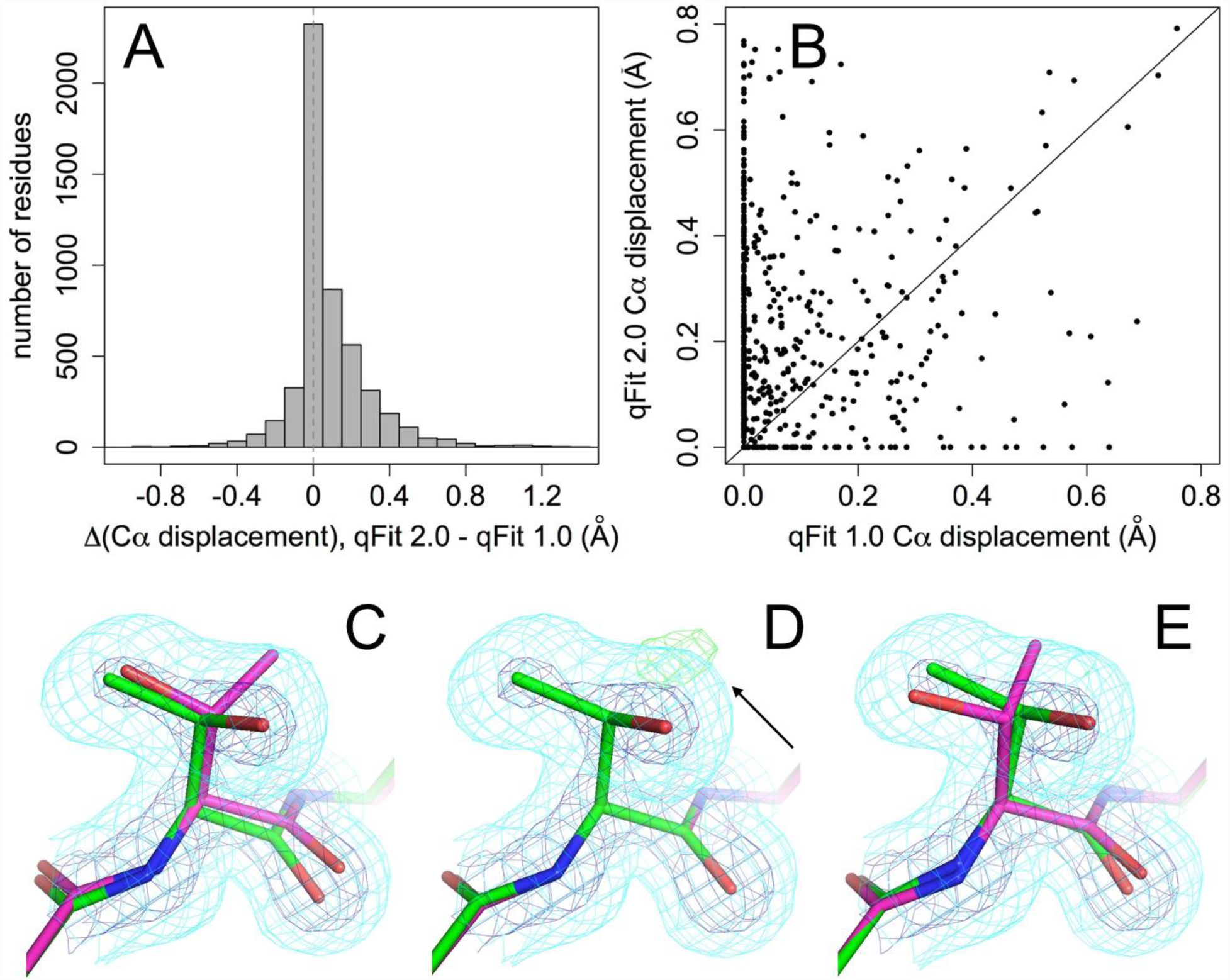
Extra backbone heterogeneity in qFit 2.0 helps discover new sidechain heterogeneity. **(A)** Histogram of difference in maximum Cα displacement across all combinations of alternative conformations between qFit 2.0 and qFit 1.0 for the test set. Vertical dotted line at 0 difference. **(B)** Maximum Cα displacements for qFit 2.0 vs. 1.0 for residues with a newly discovered sidechain rotamer in the qFit 2.0 model but not in the qFit 1.0 model. Many of these residues fall above the diagonal line, meaning the Cα moves more in the qFit 2.0 model than in the qFit 1.0 model. **(C-D)** Thr157 in cyclophilin A at room temperature (PDB ID 3k0n). 2mFo-DFc electron density is contoured at 0.5 σ (cyan) and 3.0 σ (blue); mFo-DFc difference electron density is contoured at +3.1 σ (green) and -3.1 σ (red). **(C)** The deposited structure has alternative rotamers that were correctly manually modeled. **(D)** qFit 1.0 does not move the backbone and misses the alternative rotamer, as evidenced by a peak of +mFo-DFc density (arrow). **(E)** qFit 2.0 does move the backbone (note especially the backbone carbonyl displacement), and successfully identifies the alternative rotamer.

### Newly identified peptide flips in the “flap” region of HIV protease

We also observed hidden peptide flips for the Ile50-Gly51 tight turn in the “flap” region of HIV-1 protease. HIV-1 protease is a homodimer, with residue numbers often denoted by 1-99 and 1’-99’. The flap region consisting of residues 46-56 is an antiparallel β-sheet and tight turn at the interface of the dimer (Figure 9A). In most of the hundreds of crystal structures of HIV-1 protease, the two tight turns (Leu50-Gly51 and Leu50’-Gly51’) adopt an asymmetric conformation, with one flap in a single type I conformation and the other in a single type II conformation. However, NMR relaxation data suggest that these flips can undergo chemical exchange on a slow (∼10 μs) timescale in solution [34]. Mutational data also linked collective conformational exchanges of these flips to catalytic rates [35]. In line with these solution studies, we noticed that for many HIV-1 protease crystal structures, the electron density maps actually reveal strong evidence for alternative conformations related by dual peptide flips. For example, in one high-resolution inhibitor-bound structure (PDB ID 3qih), the Leu50-Gly51 and Leu50’-Gly51’ flaps are modeled with single asymmetric conformations, but strong positive mFo-DFc electron density coincides with potentially flipped states (Figure 9B). Strikingly, qFit 2.0 automatically identifies dual “flap flips”, suggesting the flaps actually populate two different asymmetric states (green vs. purple in Figure 9C) in this particular inhibitor complex. More generally, this result suggests that these inhibitor-gating flaps in HIV-1 protease sample multiple conformations more often than previously recognized across many inhibitor complexes, which may motivate further investigation of the effects that protein and inhibitor flexibility have on binding affinity, efficiency of catalytic inhibition, and arisal of drug resistance in this biomedically important target.

**Figure 9:**
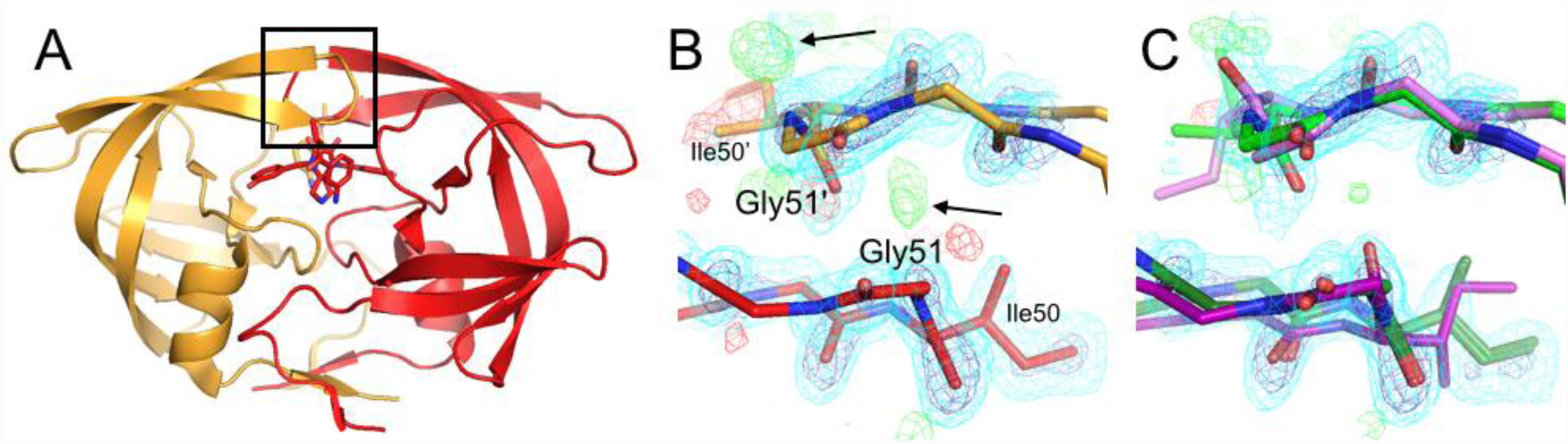
Hidden unmodeled peptide flips in the inhibitor-gating “flaps” of HIV-1 protease. **(A)** In the 1.39 Å structure of a mutant of HIV-1 protease bound to a novel inhibitor (PDB ID 3qih), the Ile50-Gly51 tight turn interacts with the dimer-related copy of itself, Ile50’-Gly51’ (boxed region). Chain A in orange, chain B in red. The inhibitor (sticks) binds in two overlapping poses immediately adjacent to these flaps. **(B)** This dimer interface, viewed as if from above in **(A)**, is asymmetric in the deposited structure: both copies of the peptide point downwards in this view. However, positive difference electron density (arrows) suggest unmodeled conformations. **(C)** qFit 2.0 models this region with coupled asymmetric peptide flips, such that both copies of the peptide point down (∼70%, green) or both point up (∼30%, purple) in this view. The multiconformer model has diminished difference electron density peaks, suggesting it is a better local fit to the data. Residual difference peaks may reflect unmodeled partial-occupancy waters that are mutually exclusive with the new protein alternative conformations. 2mFo-DFc contoured at 1.2 σ (cyan) and 3.0 σ (blue); mFo-DFc contoured at +3.0 σ (green) and -3.0 σ (red).

## Discussion

The ruggedness of protein energy landscapes leads to conformational heterogeneity even in folded globular proteins. Evidence for these alternative conformations is remarkably prevalent in high-resolution (<2 Å) crystallographic electron density maps [6]. However, because these alternative conformations are difficult and/or time-consuming to model manually using existing graphics and refinement tools, they are underrepresented in the PDB [6]. qFit is a computational approach to overcoming these problems, by automatically identifying “hidden” alternative conformations and using quadratic programming to select a parsimonious subset that collectively best explains the diffraction data. Here we have demonstrated a new version of this algorithm, called qFit 2.0, with several enhancements to handling flexible backbone -- most notably, automated detection of discrete peptide flips and explicit fitting of backbone atoms for glycines.

qFit has previously captured different types of backbone motion that can occur in secondary structure. For example, it correctly identifies the backrub motion [14] that helps Ser99 transition between sidechain rotamers in the active-site β-sheet network of CypA [15, 16], and also identifies a previously hidden α-helix winding/unwinding or “shear” motion [14, 29] (**Figure S1**). However, qFit 2.0 can now model larger backbone motions in which the backbone change itself is discrete, instead of inherently continuous but coupled to discrete sidechain rotamer changes. Specifically, it models peptide flips, which occur outside of helices and sheets and involve discrete jumps over a larger energetic barrier.

Peptide flips have important implications for understanding protein function. For example, our results for HIV-1 protease (Figure 9) strongly suggest that conformational heterogeneity, in particular peptide flips, may play underappreciated roles in protein-inhibitor complexes. Previously, molecular dynamics simulations identified a large-scale “curling” motion of these flaps that is maintained by drug-resistance mutations and therefore seems important for substrate access [36]. Although this motion is more dramatic than the peptide flaps at the tips of the flaps that we observe, it underlines that flap flexibility - potentially across multiple length scales -- is central to protease function and viral propagation. The peptide flip acts as a key conformational switch between type I/II turns, rearranging its environment beyond its immediate sequence neighbors and enabling alternative sidechain conformations with implications for function. However, the large number of unmodeled turns in HIV protease structures illustrates the challenge of distinguishing alternative conformations in electron density maps, even at high resolution. As an additional example which unfortunately lacks deposited structure factors, the active-site Gly57-Asp58 peptide in *C. beijerinckii* flavodoxin adopts distinct peptide flip states in concert with the oxidation state of the FMN prosthetic group [19]. The N137A mutation removes artificial lattice contacts that otherwise influence the conformation of the Gly57-Asp58 peptide, which results in a mixture of these peptide conformations simultaneously populated in the crystal; this suggests these multiple flip states may also coexist in solution [19].

Beyond the specific improvements to peptide flips, qFit 2.0 now fits conformations for each residue based on both sidechain (beyond Cβ) and backbone (carbonyl O) atoms. Although we originally envisioned this change for modeling glycines, we observed that it results in dramatically more extensive backbone conformational heterogeneity across the protein (Figure 8). R-factors are similar or better (Figure 4), suggesting the new models with more heterogeneity are at least as good an explanation of the experimental data. Notably, these new backbone shifts drive discovery of many more alternative sidechain rotamers (Figure 8). Our results suggest that sidechain and backbone degrees of freedom in proteins are tightly coupled, in agreement with previous reports that even subtle backbone motions can facilitate rotamer changes [14], open up breathing room for natural mutations [37], and expand accessible sequence space in computational protein design [30, 38].

Future work will investigate an armamentarium of methods for modeling larger backbone conformational change in qFit, including helix shear motions [29], adjustments of entire α-helices [39, 40], correlated β-sheet flexing [28], automated loop building algorithms such as Xpleo [9], and pre-knowledge of conformational differences between homologous structures. While these future steps will move us closer to capturing the full hierarchy of protein conformational substates [41], they will also dramatically increase the computational cost of automated multiconformer model building. Many aspects of qFit are parallelizable; however, the total computational cost for reproducing the data in this manuscript is approximately 10^5^ CPU hours. As cloud-computing capabilities of 10^8^ CPU hours can now be leveraged for pure simulation data [42], we envision that marshalling similar computational capabilities will become increasingly important for analysis of experimental X-ray data. Such data-driven computational approaches to studying the dynamic relationship between protein structure and function will be especially powerful when applied to series of datasets in which the protein is subjected to perturbations that modulate conformational distributions, such as ligand binding or temperature change [23].

## Materials and Methods

### Learning peptide flip geometries

To define possible relative geometries between flipped peptide conformations, we searched for trustworthy peptide flips modeled as alternative conformations in the Top8000 database. This database contains ∼8000 (7957) quality-filtered protein chains from high-resolution crystal structures, each with resolution < 2 Å, MolProbity score [33] < 2, nearly ideal covalent geometry, and <70% sequence identity to any other chain in the database [43 ISBN: 978-981-4449-15-1]. We searched the Top8000 for peptides with carbonyl C-O bonds pointed away from each other (O-O distance > C-C distance + 1 Å) and rotated by at least 90°, and for which both flanking Cα atoms reconverged to < 1.5 Å. Although peptide rotations of < 90° also occur, they occur more often in irregular loop regions, have less well-converged backbone for flanking residues, and are generally more diverse and difficult to simply categorize. By contrast, in this study we investigate the class of localized peptide rotations with well-converged backbone for both flanking residues. These are either very small rotations, or large flips with a rotation nearer to 180° -- the latter being the focus here. To identify test cases for qFit 2.0, we curated the resulting dataset by removing examples with more than two alternative peptide conformations; a *cis* rather than *trans* conformation for either state; or obvious errors based on steric clashes, strained covalent geometry, or torsional outliers from MolProbity [33]. This resulted in 104 examples, from which we kept a randomly selected 79 for a geometry training set (**Table S1**). We combined a subset of the remaining 25 peptide flips with a few other known examples for a test set of 18 examples (Table 1). The resolution range is 0.92-1.95 Å for the training set and 0.80-1.85 Å for the test set.

Next we characterized the geometry of peptide flips by clustering the coordinates of the flipped alternative conformation (labeled “B”) in the training set after superimposing onto a reference peptide. We used the k-means algorithm with RMSD between the five heavy atoms of the peptide backbone (Cα1, C1, O1, N2, and Cα2) for different values of k. We selected k = 4 because we observed cluster centroids with approximately 180°, +120°, and -120° rotations and for k > 4 no other significantly different rotations were identified. Notably, all four cluster centroids featured translations of the flanking Cα atoms of >0.2 Å, and as much as >0.9 Å for one cluster (“tweaked down”, red in Figure 2). The transformation matrices relating the flipped peptide cluster centroids to the reference peptide were used in qFit 2.0 to sample plausible alternative conformations, with subsequent refinement adjusting the atomic positions away from the centroid geometry.

### Tight turns and glycine enrichment

We defined tight turns as having a mainchain-mainchain hydrogen bond between *i*-1 carbonyl C=O and *i*+2 amide N-H that was detectable by the program Probe [44]. This definition is somewhat conservative; several more examples also were visually similar to tight turns. Enrichment of glycines at the two positions involved in a peptide flip was assessed for different peptide flip clusters within the training set relative to a large set of 337 randomly selected structures containing 6,092 total glycines out of 78,094 total amino acid residues. The statistical significance of this enrichment was assessed using a one-tailed Fisher’s exact test based on the hypergeometric distribution [45].

### qFit

#### qFit part 1: Preparing each residue for qFit

qFit exhaustively examines a vast number of interpretations of local electron density, and deterministically selects a small ensemble that optimally explains the density. The method starts from an initial single-conformer model. The occupancies of all atoms in a residue, *k*, beyond the Cβ atom are set to zero with phenix.pdbtools, and the model is refined with phenix.refine. Refinement uses anisotropic B-factors for all residues if the resolution is better than 1.45 Å, or just for residue *k* otherwise. Finally, all atoms in residue *k* beyond the Cβ atom are removed. These steps result in two inputs to qFit: (1) an omit map and (2) starting coordinates with an anisotropic tensor for the Cβ atom.

#### qFit part 2: Peptide flips and backbone sampling

Next, the peptide from residue *k* to *k*+1 is aligned to the centroids identified from clustering the Top8000 dataset (see above). We calculate local coordinate frames for the peptide and cluster centers by orthogonalizing the Cα_i_-Cα_i+1_ and Cα_i_-O_i_ vectors and taking their cross-product. Each centroid conformation is then transformed onto the starting peptide using a homogeneous coordinate transformation, resulting in a candidate flipped alternative conformation.

Peptide flips do not occur in canonical secondary structure due to steric constraints, so qFit 2.0 does not attempt them in helices and sheets, as detected by the CCP4 MMDB library [46]. This both avoids false positives and affords a computational speedup by reducing combinatorics in the selection steps (see below).

Next, for each residue *k*, a fragment of length 7 centered on residue *k* is extracted. For each candidate conformation (one unflipped plus four flipped), the Cβ atom is moved along the major and minor axes of the ellipsoid (six total directions) by a distance determined by the ellipsoid eigenvectors and a scale value provided by the user. Here we used 0.1, 0.2, and 0.3 for this scale value, and 0.05 for the optional value for random additions to scale. For glycines, which lack a Cβ atom, the backbone O atom is used to define the anisotropic ellipsoid. To preserve the exact geometry of the fragment, we use inverse kinematics to deform the fragment. The gradient of the distance function is projected onto the nullspace spanned by the dihedral degrees of freedom of the fragment [9, 10]. These motions further position backbone atoms to accommodate rotameric sidechain conformations.

#### qFit part 3: Sidechain sampling

For small sidechains (Ala, Asn, Asp, Cys, Gly, Iso, Leu, Pro, Ser, Thr, Val), a 40° neighborhood of each rotameric X dihedral angle, starting at -20°, is sampled in 10° increments on each of the 35 backbone conformations. To avoid a combinatorial explosion, large sidechains (Arg, Glu, Gln, His, Lys, Met, Phe, Trp, Tyr) are sampled hierarchically. First, the backbone and first dihedral angle are sampled similarly to small sidechains. A larger neighborhood of 50° is sampled in 4.5° increments to avoid missing conformations that are initially suboptimal but can accommodate better fits for subsequent X angles. This set is then subjected to the selection procedure, which returns a handful of conformations that fit the density up to the Cγ atom. These selected conformations form the basis for sampling the next X angle using the same parameters. This procedure is repeated until the entire sidechain is built.

#### qFit part 4: Conformer selection

For each of the *N* conformations sampled for each residue, we calculate an electron density map 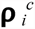. We scale the observed electron density map ***ρ**^o^* to ***ρ**^c^*. We then subject the weighted sum of 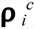 to a quadratic program (QP) to determine a vector of occupancies *w*^*T*^ that minimizes the least squares residuals with respect to the observed electron density:

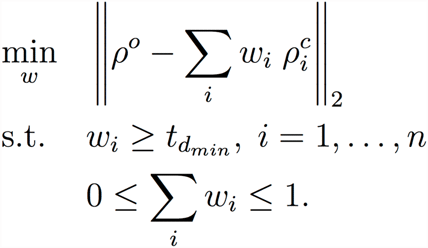

The residuals are calculated over regularly spaced voxels that are within a resolution-dependent radius *r* of any of the sidechain (Cβ and beyond) or carbonyl O atoms. The radius *r* (in Å) is determined by *r* = 0.7 + (*d* - 0.6)/3.0 if d < 3.0Å, and 0.5 d if d ≥ 3.0Å, where *d* is the resolution in Å.

The vast number of conformations results in a system that is generally underdetermined. We therefore enforce sparsity of the solution by introducing a resolution-dependent *threshold* constraint 0 < *t*_*dmin*_ *≤ 1* for the occupancies; i.e., *w*_*i*_ > *t*_*dmin*_ for all *i*. The threshold constraint prevents overfitting by suppressing arbitrarily small occupancies that model noise. Together with the constraint that the total occupancy cannot exceed unity, the threshold also enforces a *cardinality* constraint; i.e., the number of non-zero occupancies is bounded by the integer part of 1/*t*_*dmin*_. In effect, the threshold constraint enforces selection of an optimal subset in the regression. Note that the two constraints imply *w*_*i*_ ∈ {0} ∪ [*t*_*dmin*_,1]. Introducing binary variables *z*_*i*_ ∈ {0,1}, we can rewrite the optimization problem as a mixed integer quadratic program (MIQP):

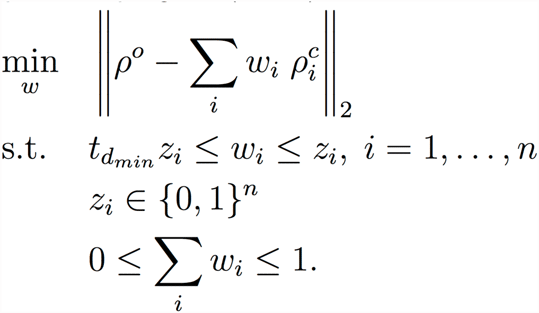

This optimization problem belongs to the class of convex quadratic problems, for which solvers can find a globally optimal solution. An MIQP is NP-hard. We therefore pre-fit conformers with QP, and subject all conformations with non-vanishing occupancies to MIQP. While in theory this no longer guarantees an optimal solution, practice tests on small sets of conformers did not show an effect of pre-fitting.

#### qFit part 5: Putting the model back together

Assembling a consistent, multiconformer model from individually fitted residues requires two steps. First, backbone heterogeneity can extend over multiple, consecutive residues, each with slightly different occupancies and/or numbers of conformers. To synchronize the number of conformers and their occupancies over a fragment of length *K* residues consisting of consecutive backbone multiconformers, we again rely on conformational selection by MIQP. We enumerate all possible connections between all conformations *C*_*i*_ of residues *i = 1, …, K* to obtain *C*_*f*_ = ∏ ^*K*^_*i=1*_*C*_*i*_ conformations to model this fragment. We subject all *C*_*f*_ conformations to the MIQP, which selects a parsimonious ensemble of at most 1/*t*_*dmin*_ conformations based on optimal fit to the observed electron density, each with identical occupancy for all atoms in the fragment. Note that *C*_*f*_ can be quite large, even for modestly long fragments. To avoid a combinatorial explosion, for long fragments we implemented a divide-and-conquer procedure that fits segments of each fragment with MIQP. The fitted segments are then combinatorially recombined and again subjected to MIQP to obtain the final set of conformations for the fragment. The peptide-bond geometry of the output model at this stage of qFit can be distorted. A later refinement stage with phenix.refine corrects the geometry.

Second, conformations encoding collective motions are often mutually exclusive. In crystal structures, each internally consistent set of residues is labeled with an alternative conformation or “altloc” identifier -- a capitalized letter (“A”, “B”, etc.) for multiple conformations or a blank space (“ “) for a single conformation. However, the initial model from the preceding steps in the qFit pipeline has random labels. To identify internally consistent labels, we use a simple downhill Monte Carlo optimization protocol. The program Label minimizes a simple Lennard-Jones score by randomly swapping labels between conformations and accepting the change if the score improves. This is repeated 10,000 times per trial over 10 trials, and the model with the best score is used for subsequent steps.

To finalize the model, we first refine the relabeled model with phenix.refine. Next, we remove conformations that are now indistinguishable from other conformations within predicted coordinate error, and reset occupancies to sum to unity for each atom. Finally, we refine again, using anisotropic B-factors if the resolution is better than 1.45 Å.

### Hydrogen treatment

Hydrogens were placed at nuclear positions for Label in qFit 1.0 and at electron-cloud positions for Label in qFit 2.0. Correspondingly, for Label in qFit 2.0, hydrogen van der Waals radii were taken from the new values in Reduce [47], which are intended to match those used in PHENIX. Hydrogens were absent for all other steps in qFit, including the final refinement step; however, the user is encouraged to add hydrogens to the final qFit model for their protein of interest and proceed to other analyses. Future work will update programs for downstream analysis of qFit models such as CONTACT [16] to also use electron-cloud instead of nuclear hydrogen positions.

### Generating synthetic datasets

To generate synthetic datasets for testing qFit, we used the protein chains containing the four peptide flip cluster centroids (3mcw B 101-102, 2ior A 159-160, 2g1u A 51-52, 3g6k F 172-173). We first used phenix.pdbtools to convert any anisotropic B-factors to isotropic, added 10 Å^2^ to each B-factor per Å of resolution worse than the original structure’s resolution to roughly simulate the general rise of B-factors with resolution, and placed the chain in a P1 box that comfortably encompassed it. Next we used phenix.fmodel to calculate structure factors (with the “k_sol=0.4” and “b_sol=45” bulk solvent parameters, and also generating 5% R-free flags) and added 10% noise in complex space with the sftools utility in CCP4 [46]. This process was repeated for every simulated resolution from 0.9 to 2.0 Å with a 0.1 Å step size.

### Evaluating true and false positives

qFit uses an input parameter (MC_AMPL) to scale the magnitude of movements of the Cβ (or O for glycines) along the directions dictated by its thermal ellipsoid. As in previous work [10, 16, 26], we explored multiple values for this parameter: 0.1, 0.2, and 0.3. For evaluating results such as true vs. false positive peptide flips and rotamers here, we considered all three resulting qFit models for each dataset. This is sensible because an end user of qFit 2.0 will likely reproduce this same protocol (with a few MC_AMPL values) and thus have a choice of models to use for developing insights into conformational heterogeneity and its connection to function. For other analyses, we used the minimum-R_free_ qFit model model unless otherwise noted.

### Re-refinement with water picking

To compare R-factors between the deposited models and qFit 2.0, we finalized both models with phenix.refine for 10 macro-cycles using the same parameters, including the “ordered_solvent=true” flag. The resulting R-factors for qFit 2.0 models are similar or slightly better (Figure 4).

### Programs and databases

PHENIX version 1.9-1692 (the most recent official release) [48] was used for all steps of both qFit 1.0 and 2.0. Coordinates and structures factors were obtained from the Protein Data Bank [49]. qFit uses the following libraries: IBM’s ILOG CPLEX solver for QP and MIQP, which is available free of charge for academic use, and LoopTK for inverse kinematics calculations [50]. qFit is implemented in parallel; it is capable of sampling and evaluating conformations for each residue as an independent job on a Linux cluster. We have implemented job management for qFit on both Oracle/Sun Grid Engine and LSF Platform.

## Supplementary Figures

**Figure S1:**
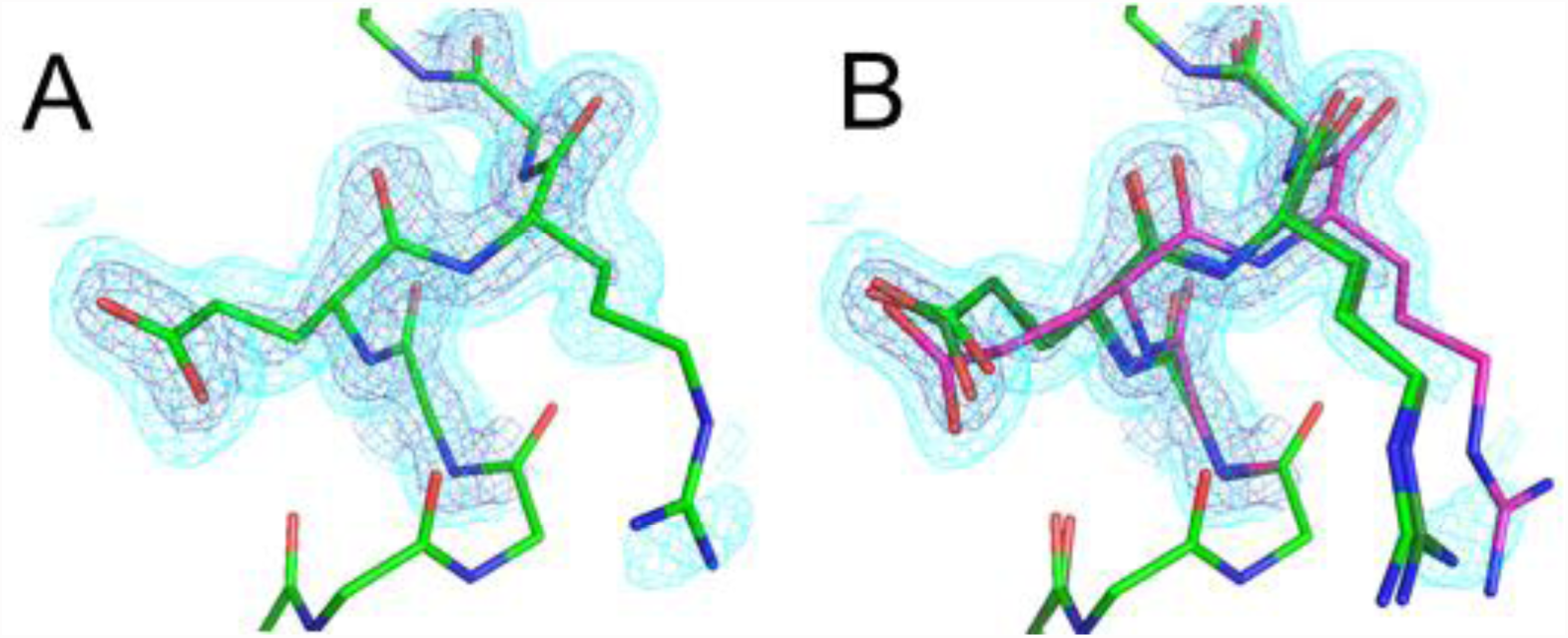
qFit detects a shear backbone motion in a room-temperature crystal structure of cyclophilin A. **(A)** Residues 142-145 in CypA are modeled with a single conformation in the single-conformer structure (PDB ID 3k0n). The model is a reasonable fit to the 2mFo-DFc electron density contoured at 1.0 σ (cyan) and 2.5 σ (dark blue), which is slightly anisotropic for the central carbonyl oxygen. **(B)** The multiconformer qFit model, on the other hand, includes three alternative conformations with backbones related by a shear-like motion to explain the electron density. Each shear end-state (greens vs. purple) is allocated about 50% occupancy. The multiconformer model adds a second rotamer (purple) in addition to the original rotamer (greens) for Glu143 (left-hand-side of panel) and sweeps the Arg144 sidechain sideways (right-hand-side of panel).

**Figure S2:**
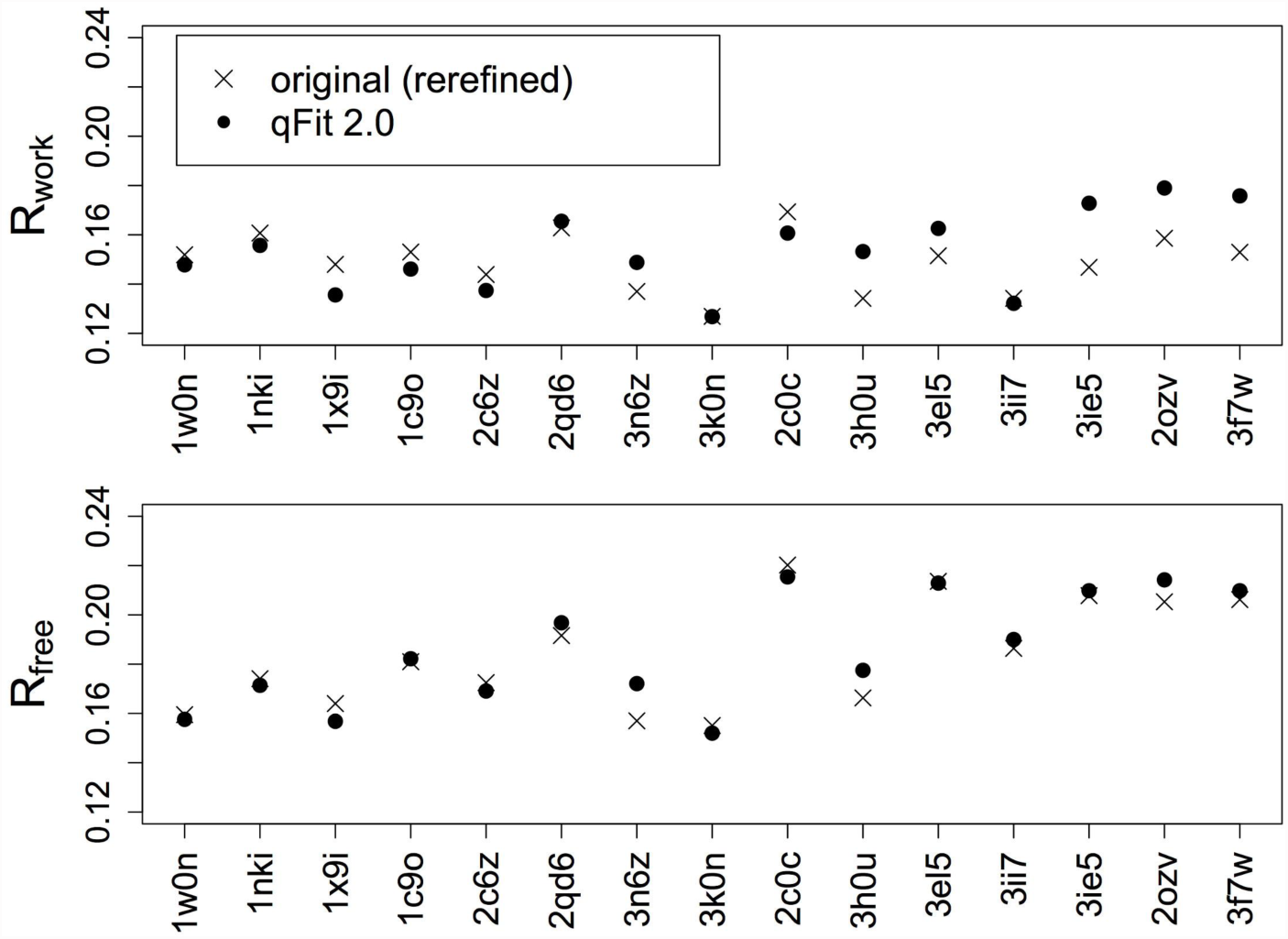
Multiconformer modeling with qFit does not result in better crystallographic R-factors before solvent picking. R_work_ and R_free_ are plotted vs. PDB ID sorted from high to low resolution. X’s indicate original structures rerefined without automated addition and removal of water molecules, and filled circles indicate qFit 2.0 models.

**Table S1:** List of peptide flip examples from Top8000 used as training set.

